# Preclinical translation of *Neurofibromatosis type 1* (*NF1)* exon 17 skipping using targeted U7-SnRNA packaged into engineered AAV serotypes

**DOI:** 10.64898/2026.06.29.734312

**Authors:** Marc Moore, Kimia Rayat-Sanati, Xiaoxia Zhang, Hui Liu, Zakaria Rostamitehrani, Tarah Vijayasarathy, Erik Westin, Miguel Sena-Esteves, Casey A. Maguire, Robert A. Kesterson, Linda Popplewell, Deeann Wallis

**Author notes:** Corresponding author: Department of Genetics, Heersink School of Medicine, University of Alabama at Birmingham, Kaul Rm 640A, 720 20th St South, Birmingham AL 35294, |.

## Abstract

To facilitate the translation of *NF1* exon 17 skipping as a mutation-specific therapy for Neurofibromatosis type 1 into *in vivo* testing, we have continued to develop more efficient antisense oligonucleotides (ASOs), humanized mouse models, and explored multiple delivery platforms including an adeno-associated virus (AAV)-U7-SnRNA vector approach. We evaluated both biodistribution and exon skipping efficacy of a U7-SnRNA targeting *NF1* exon 17 with an SFFV-driven cassette containing T2A-linked Luciferase (Luc) and eGFP packaged in AAV-9, AAV-F and AAV-B1 capsids. We show that AAV-F is superior to AAV-9 and AAV-B1 for mouse brain delivery based on DNA transduction, GFP expression, and luciferase activity, but AAV-B1 delivers 2-4 fold more to sciatic nerve (SCN). In terms of exon skipping, AAV-F appears to induce the most skipping in liver and optic nerve (ON), while AAV-B1 mediates highest skipping in the liver, SCN, and ON. The identification of AAV serotypes that allow efficient transduction and delivery of transgenes to the mouse CNS and PNS is impactful for preclinical research in murine models of other diseases. Furthermore, this is both the first report of *NF1* exon skipping efficacy *in vivo* and the first successful application of an U7-SnRNA for the restoration of functional neurofibromin for NF1.

**Graphical Abstract:** 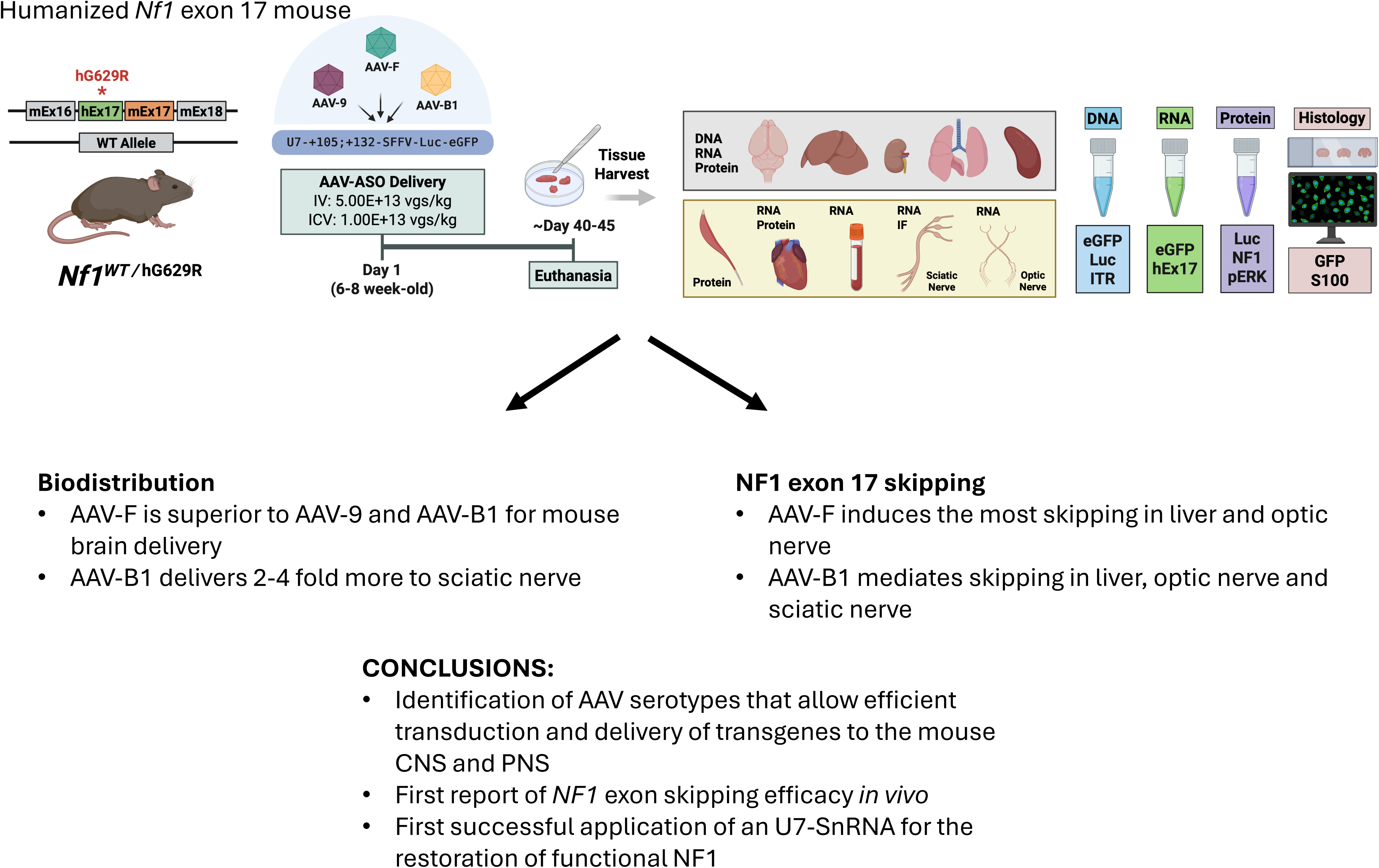

## INTRODUCTION

Neurofibromatosis Type 1 (NF1) is a tumor predisposition disorder caused by pathogenic variants in the *NF1* gene. Neurofibromin, the protein encoded by *NF1*, functions as a negative regulator of RAS signaling and thus acts as a tumor suppressor. Estimates indicate that NF1 affects 1 in 2,500-3,000 individuals worldwide^1–3^. This prevalent monogenic disease has multi-system effects and pleomorphic phenotypes across the central and peripheral nervous systems^4^. Early NF1 typically presents with café au lait macules (CALMs), ocular Lisch nodules, and skeletal abnormalities. Subsequent acquisition of additional pathogenic variants in the second *NF1* allele results in hyperactivation of the RAS/MAPK pathway, uncontrolled cellular proliferation, and the development of NF1-associated tumors^5^. Some affect the CNS, as in the case of optic pathway gliomas and low-grade brain tumors; whereas cutaneous neurofibromas (cNFs) and plexiform neurofibromas (pNFs) affect the PNS and arise from superficial and deep nerve structures, respectively. Whilst cNFs are usually benign, approximately 8-13% of pNFs can undergo malignant transformation into malignant peripheral nerve sheath tumors (MPNSTs), a process associated with poor clinical outcome^6^. Crucially, NF1 patients are 1.8 times more likely to develop other cancerous malignancies, and estimates suggest a 15–20-year reduction in life span relative to the general population^7^.

Currently, clinical interventions for this severe condition are limited and focus on tumor surveillance and palliative care. Surgical resection of tumors is the primary intervention. However, the invasive nature of the tumor can make complete resection challenging while avoiding nerve and proximal physiological structures, and thus, tumor recurrence often occurs. In specific cases, radiation and chemotherapy are used as adjuvant therapies. Recently, two MEK inhibitors, Selumetinib and mirdametinib, were approved by the FDA to treat pNFs, but neither drug prevents malignant transformation. In addition, these drugs require long-term dosing to suppress tumor growth but have numerous side effects, including dermatological toxicity and gastrointestinal disturbances, resulting in low treatment compliance^8–10^. In addition, MEK inhibitors do not work for 30% of patients with symptomatic inoperable pNFs, and for those amenable to treatment, resistance can develop. There is a clear unmet clinical need for more comprehensive therapeutic options addressing NF1-associated morbidities and mortality.

Extensive research of advanced therapeutics to more adequately address the needs of NF1 patients is ongoing. Development of classical gene replacement therapies has been slow, hindered by the 8.5kb size of the *NF1* cDNA, and over 3100 pathogenic variants being reported across the gene, without a focal mutational hotspot. Importantly, the full *NF1* cDNA exceeds the ∼4.5kb packaging capacity of Adeno-Associated viruses (AAVs), the favored gene therapy vector due to high efficiency of *in vivo* gene transfer and permanence of transgene expression. To provide a therapeutic strategy within the limited transgene capacity of AAV, several miniNF1 protein configurations carrying the GAP-related domain (GRD) alone or fused to membrane-localization domains have been shown to regulate Ras pathway activity and control tumor growth *in vivo*^11, 12^. Despite promising results in animal models, these miniNF1 proteins are unlikely to replace all functions of the full-length NF1 protein. AAV-mediated gene therapy approaches are also further complicated by limited transduction and unregulated expression level of encoded transgenes. Naturally occurring AAV serotypes have shown limited transduction of the central and peripheral nervous systems, prompting extensive AAV capsid engineering efforts to achieve more efficient and specific transduction profiles^13–15^. AAV-F, an engineered AAV-9 variant identified by *in vivo* capsid library selection, enables over an order of magnitude higher crossing of the blood-brain barrier in mice than AAV-9 after IV infusion^16^. This is driven by the Ly6c1 receptor on murine brain endothelium^17^. In subsequent studies, AAV-F was shown to robustly transduce gray matter of the spinal cord after lumbar intrathecal delivery in adult mice in comparison to AAV-9^13, 16^. AAV-B1, generated through parental capsid shuffling of multiple AAV serotypes, robustly transduces the mouse brain, spinal cord, muscle, pancreas, and lung after IV infusion in adult mice^14^. One of the major challenges of AAV gene therapy for neurological diseases, in the context of disease relevant tissue, is the heterogeneous transduction across cellular populations. This heterogene leads to variability in intracellular vector genome copy number and limits precise control of transgene expression levels and can contribute to dose-related toxicity^18^. One approach to control this is the judicious selection of neurological promoters and synthetic promoters with capacity for homeostatic maintenance.

In contrast to AAV-based gene augmentation approaches, antisense oligonucleotide (ASO) mediated exon-skipping aims to modulate splicing at the pre-mRNA level by targeting splice acceptors/donors, cryptic splice sites, or exonic splicing enhancers. This approach can be employed to mask exons with pathogenic variants, thereby restoring the reading frame and/or protein expression/function, albeit with a small internal deletion^19^. Exon skipping using phosphorodiamidate morpholino oligomer (PMO) sequences have clinical precedence as seen in Duchenne Muscular Dystrophy (DMD)^20^, with FDA approval granted for Exon 45 (Amondys)^21^, 51 (Exondys)^22, 23^ and 53 (Vyondys) skipping^24^. Critically, in exon-skipping, gene expression remains under control of the endogenous promoter, allowing for proper physiological responses. In addition, exon skipping ASOs produce a truncated protein with closer structure and functionality to the full-length. Finally, as this is an established therapeutic modality, numerous studies have sought to address challenges of oligo-related delivery via conjugation to peptides or vectorization. Importantly, this provides a wider toolbox for successful therapeutic translation.

Interestingly, there has been limited exploration of exon skipping for NF1. Early studies focused on masking deep intronic cryptic splice sites (CSS) to prevent inclusion of intron sequences in the *NF1* mRNA. Two studies demonstrated that ASOs can effectively mask CSS and restore normal splicing *in vitro*, resulting in an antisense-dependent decrease in RAS-GTP levels^25, 26^. An inherent advantage of such approaches is that the full-length *NF1* mRNA is restored. However, there is little research on targeting ASOs to pathogenic sequences in *NF1* exons. We previously investigated which exon coding sequences would be amenable to exon skipping without significant loss of neurofibromin expression or function^27^. Exon 17, amongst others, was identified as a candidate. Our *in vitro* work demonstrated that treatment of Schwann cells with ASOs targeting exon 17 restored neurofibromin expression and reduced RAS pathway activity. Importantly, mouse models of Exon 17 deletion that are missing exon 17 on both alleles also showed no tumorigenesis indicating therapeutic promise^27^.

Translation of exon skipping into *in vivo* models is challenging due to limited humanization of mouse models and ASO delivery challenges. In this work, we further refined ASOs targeting *NF1* exon 17, encoded antisense sequences into AAV-U7-SnRNA constructs, and developed a new mouse model with a humanized *NF1* Exon 17. In parallel, we developed an AAV-U7-SnRNA vector encoding a Luciferase-eGFP bicistronic reporter to assess gene transfer efficiency using AAV-9 and two additional capsids, AAV-F and AAV-B1, engineered to have enhanced CNS/PNS tropism, to determine whether more efficient gene transfer improves exon-skipping efficacy. Here we provide the first evidence of *in vivo NF1* exon skipping that restores functional neurofibromin expression. In addition, from comparison of intracerebroventricular (ICV) and intravenous (IV) routes of administration, we demonstrate that AAV-F has superior targeting to brain relative to AAV-9 and AAV-B1 in mouse models. AAV-B1 shows an increased propensity for SCN transduction and exon skipping, indicating AAV capsid selection for mouse model testing may be influenced by disease indication.

## MATERIALS and METHODS

### Development of ASOs

Developing upon our prior work, PMO designs were refined^27^. The *NF1* Exon 17 sequence was input into sOligo software (https://sfold.wadsworth.org/cgi-bin/soligo.pl) keeping default folding temperature of 37°C and ionic conditions of 1M NaCl with no divalent ions to ascertain an *in silico* prediction of secondary RNA structure.

PMOs were designed using sFold (http://www.unafold.org/mfold/applications/rna-folding-form.php.) The output PMOs were selected based on the number of exonic splicing enhancer (ESE) motifs masked, their Gibbs free energy ≥-8.0 and the mapping of these PMOs to “open” and “closed” regions of the secondary structure

Furthermore, positional clustering to PMOs with known efficacy was also considered. Notably, PMO “+108; +132” previously disclosed retained on-target activity in HEK293T/A15 cell lines, HEK293T cell lines engineered to have c.1885G>A: pG692R frameshift mutation in Exon 17 of the *NF1* gene. Thus, a tiled array around this position was undertaken to find higher efficacy candidates. From this screen we resolved upon +105; +132 as the lead candidate in this investigation.

### Development of pAAV-U7-SnRNA Constructs

To enable the vectorization of antisense oligonucleotide sequences, a gBlock containing a U7-SnRNA transgene was synthesized. This comprised the U7 promoter sequence, a stuffer sequence flanked by Type IIS endonucleases, an SmOPT sequence, a U7 hairpin and a downstream tail, analogous to published sequences^28^. Notably, this gBlock was constructed to have multiple cloning sites on the 5’ and 3’ end compatible with the pAAV-MCS plasmid. The gBlock was then sub-cloned into the pAAV-MCS plasmid, using SphI and NdeI restriction endonucleases. Sanger sequencing was then used to confirm the identity of the insert. Once confirmed, constructs were maxi-prepped to enable the sub-cloning of desired antisense sequences.

Antisense sequences were synthesized as two compatible primers with overhangs corresponding with the type IIS restriction sites and hybridized in Roche H- buffer. The use of the type IIS restriction enables the antisense sequences to be sub-cloned as contiguous with the U7-transgene.

### AAV Viral Preparation

Recombinant AAV vectors were produced by a double-transfection plasmid system in the case of AAV-9 comprising of the gene of Interest: pAAV-U7-SnRNA / pAAV-U7-SnRNA-SFFV-Luciferase-T2A-eGFP and the pAAV helper Cap9 (pDF9 helper plasmid encoding viral cap,rep ORFs and Adenoviral helper genes).

Alternatively in the case of AAV-B1 and AAV-F a triple transfection system was utilized comprising of the gene of Interest: pAAV-U7-SnRNA / pAAV-U7-SnRNA-SFFV-Luciferase-T2A-eGFP, adenovirus helper plasmid: pAdDeltaF6 (University of Pennsylvania; Addgene #112867) or pAldX80 from Aldevron and the capsid plasmids: AAV-F (AAV-F in pAR-9 was a gift from Casey Maguire (Addgene plasmid # 166921; http://n2t.net/addgene:166921; RRID:Addgene_166921)) or AAV-B1 (Umass Chan Medical School; Addgene #78504)^16^.

All transfections were performed in HEK293T/C17 (ATCC) using a 1:1 molarity of plasmids and Polyethyleneimine Max (PEI Max, MW40,000 Polysciences, #24765) in Dulbecco’s modified Eagle’s medium with 2% fetal calf serum. Three days later cells were lysed and recombinant AAV clarified and purified with HPLC using POROS CaptureSelect AAV-9 resin (Thermo Fisher Scientific). Once eluted the AAV was dialyzed and resuspended in PBS-MK (phosphate buffered saline with 5mM MgCl2 and 12.5mM KCl). Using Millipore Amicon Ultra 15 (100k MWCO) spin columns, the AAVs were concentrated and quantified by quantitative PCR (qPCR).

### Development and characterization of *Nf1* humanized exon 17 with G629R insertion mouse. CRISPR/sgRNA design and synthesis

CRISPR guides were designed to humanize an extra 50 bp for each intron using CRISPOR (crispor.tefor.org) (5’ = acactgacagagtgcttagt ggg; 3’ = ttaccaacatccactcagaa agg). Modified synthetic sgRNAs (Synthego, Inc.) were allowed to complex with Alt-R® S.p. Cas9 Nuclease V3 (IDT, Inc.) at room temperature for 15 minutes before dilution with DMEM to a final concentration of 100 ng/ul, 100 ng/ul and 200 ng/ul of guide 1, guide 2 and Cas9, respectively. We also included a repair template (TGAGTGCTTCTAACGGGAAAGCTTTTGGAGAGAAGGTTTTCCAGGGCTGCCTTAAAATAGAAAGTCAGTTCCCACCTCTTGGTTGTCAGTGCTTCAGTAAAGCTTATTTATTTATTTTTTTCTAGCAGGCAGATAGAAGTTCCTGTCACTTTCTCCTTTTTTACaGGGTAGGATGTGATATTCCTTCTAGTGGAAATACCAGTCAAATGTCCATGGATCATGAAGAATTACTACGTACTCCTGGAGCCTCTCTCCGGAAGGGAAAAGGGAACTCCTCTATGGTCAGCTTCTTCTGTACTTTTTCTGTATCATTTTATGTGCTCTGTTTGTTTTCTGAGTGGATGTTGGTAAATTTCGATTAAATAATTTAGCACATTAGTTACAGACTGCTCAGAATTTAGTTGTA).

### Gonadotropins

Female C57B6J embryo donors from 3-4 weeks of age were administered 5 IU of PMSG (Sigma, St. Louis, MO, USA) on day -3 followed by 5 IU of HCG (Sigma, St. Louis, MO, USA) on day -1 to induce superovulation. Donor and recipient females were mated to stud and vasectomized males, respectively, on day -1.

### Collection of embryos

At day 0.5 post-conception, super ovulated donor females with copulatory plugs were humanely sacrificed using CO_2_ followed by cervical dislocation. Oviducts were dissected into sterile medium and nicked to expose the cumulus masses containing fertilized embryos. Embryos were cultured in KSOM (Millipore, Darmstadt, Germany) under 5% blood-gas prior to electroporation.

### Electroporation

Fertilized embryos were placed in a 5 mm petridish parallel platinum plate electrode (Nepa Gene, Co., Chiba, Japan) and electroporated using the NEPA21 Super Electoporator in batches of 25 using the following parameters: Poring 150V, 3 ms, No. 4, Decay Rate 10%, Polarity +; Transfer 20V, 50 ms, No. 5; Decay Rate 40%, Polarity +/-. Surviving embryos were transferred as previously described.

### Animal identification

Animals were identified by cage card, sex, and unique ear tags consecutively numbered that were affixed at weaning.

### Biopsies

Tail biopsies were collected at weaning. A 5-7 mm portion of the distal segment of the tail was cut for analysis and the remainder cauterized. Genomic DNA was purified from the lysed tail samples.

### Identification of founders

Chimeric pups were genotyped and we identified at least one putative founder that may recapitulate the variant of interest and be useful for testing the ASOs for exon skipping. Founder animals were identified by PCR using primers flanking the target loci: F AAGTTCTGGGGAACAGGAACG and R GGTCAGCCACACGGTCAAAT (hEx17-LgAmpl-1 primers). This created two amplicons, one 621 bp wildtype and one ∼1800bp. We gel purified and Sanger sequenced bands. Sequencing indicated that instead of replacing mouse exon 17 with humanized exon 17, there was a 5’ insertion of the entire humanized exon 17 with the desired patient mutation along with 28 bp of human intron 16 5’ of the humanized exon 17 and 50 bp of human intron 17 3’. Mouse exon 17 is 41 base pairs downstream of the engineered locus.

### Tissue Preparation and Isolation of DNA and RNA

Tissue samples (∼10 mg) were homogenized using a bead-based homogenizer in the appropriate 350ul Buffer RLT. Genomic DNA and total RNA were simultaneously isolated using the AllPrep DNA/RNA Mini Kit (Qiagen, Cat# 80204) according to the manufacturer’s instructions. Briefly, lysates were applied to AllPrep DNA spin columns to isolate genomic DNA, followed by RNA purification from the flow-through using RNeasy spin columns.

On-column DNase digestion was performed using the RNase-Free DNase Set (Qiagen, Cat# 79254) to eliminate genomic DNA contamination. Purified RNA was eluted in RNase-free water, and DNA was eluted in EB buffer. Nucleic acid concentration and purity were assessed using a NanoDrop 2000 spectrophotometer (Thermo Fisher Scientific).

### Quantitative PCR for Genomic DNA

Quantitative PCR for genomic DNA (DNA qPCR) was performed using Luna Universal qPCR Master Mix (New England Biolabs, Cat# M3003). Each 10 µL reaction contained 5 µL 2× master mix, 0.25 µM forward and reverse primers, and 10 ng genomic DNA template. Primers: eGFP F: GAACCGCATCGAGCTGAA, eGFP R: TGCTTGTCGGCCATGATATAG; Inverted Terminal Repeat (ITR) F: GGAACCCCTAGTGATGGAGTT, ITR R: CGGCCTCAGTGAGCGA; Luciferase (Luc) F: ATGGACAGCAAGACCGATTAC, Luc R: GCTTGAAGTCGTACTCGTTGA

Thermal cycling conditions were as follows: initial denaturation at 95°C for 1 min, followed by 40 cycles of 95°C for 15 s and 60°C for 30 s. Standard curves were generated using serial dilutions of AAV-9 standards.

### Reverse Transcription and Quantitative PCR (RT-qPCR)

Reverse transcription and quantitative PCR were performed using the Luna Universal One-Step RT-qPCR Kit (New England Biolabs, Cat# E3005). Each 10 µL reaction contained 5 µL 2× Luna Universal One-Step Reaction Mix, 0.5 µL Luna WarmStart RT Enzyme Mix, 0.4 µM forward and reverse primers, and RNA template 10 ng.

Thermal cycling conditions were: reverse transcription at 55°C for 10 min, initial denaturation at 95°C for 1 min, followed by 40 cycles of 95°C for 10 s and 60°C for 30 s. Melt curve analysis was performed to verify amplification specificity. Primer: eGFP F: GAACCGCATCGAGCTGAA, eGFP R: TGCTTGTCGGCCATGATATAG. mEX17 F: TGGGAAATCAGCTCACAAATGC, hEx17 R: CCTTCCGGAGAGAGGCT, mEX15 F: CTGGGAAATCAGCTCACAAATG, mEX16 R: TCCCGTAACCACTTGAGAATTT.

Relative gene expression levels were calculated using the 2^(-ΔΔCt) method with Actin as the internal control.

### Western Blot Analysis

Protein extracts were prepared in RIPA buffer supplemented with protease inhibitors (Roche, Cat# 05892791001) and PhosSTOP™ (Roche, Cat# 04906837001). Protein concentration was determined using the Pierce BCA Protein Assay Kit (Thermo Fisher Scientific, Cat# 23227). Equal amounts of protein (20–50 µg) were resolved by SDS-PAGE and transferred onto PVDF membranes (Bio-rad Cat# 1620177). Membranes were blocked with 5% BSA in TBST for 1 h at room temperature and incubated overnight at 4°C with primary antibodies. Antibody: NF1 (abcam, cat# ab17963 1:1000), b-actin (Cell Signaling cat# 3700 1:1000), p-ERK (Cell Signaling cat# 9101 1:1000), and total ERK (Cell Signaling cat# 9102 1:1000). After washing, membranes were incubated with HRP-conjugated secondary antibodies for 1 h at room temperature. Signals were detected using Clarity Western ECL Substrate (Bio-rad, Cat# 170-5061) and visualized using a ChemiDoc MP Imaging System (Bio-Rad). Band intensities were quantified using ImageJ software and normalized to β-actin.

### Luciferase Reporter Assay

Luciferase activity was measured using the ONE-Glo™ Luciferase Assay System (Promega, Cat# E6120) according to the manufacturer’s instructions. Briefly, tissue samples expressing luciferase were lysed in RIPA lysis buffer with protease inhibitors. Lysates were clarified by centrifugation at 12,000 × g for 15 min at 4°C, and the supernatants were collected for analysis. Total protein concentration was determined using the Pierce BCA Protein Assay Kit (Thermo Fisher Scientific, Cat# 23227) to allow normalization of luciferase activity to protein input. Equal amounts of protein (typically 30 µg in 30 µL volume) were transferred to white opaque 384-well plates. ONE-Glo™ Luciferase Assay Reagent was equilibrated to room temperature and added to each sample at a 1:1 ratio, 30 µL lysate + 30 µL reagent. Plates were gently mixed for 1–2 min and incubated at room temperature for 3 min to allow complete reaction development. At the indicated time point, the ONE-Glo™ reagent was equilibrated to room temperature and added directly to each well at a 1:1 ratio with the culture medium (e.g., 100 µL reagent to 100 µL medium). Plates were gently mixed for 2 min to ensure complete cell lysis and incubated at room temperature for 3 min to stabilize the luminescent signal. Luminescence was measured using a microplate luminometer. Background signal from lysis buffer plus reagent controls was subtracted from all measurements. All samples were analyzed in technical triplicates with at least three independent biological replicates.

### Histology

Harvested tissues were fixed in 4% PFA at 4°C overnight and then placed in 30% sucrose at 4°C until the tissue sinks. Tissues were then embedded in OCT and rapidly frozen directly on dry ice and stored at −80 °C. Tissues were cut it into 10 µm sections and mounted onto microscope slides.

### Immunofluorescence staining

Slides were washed twice with PBS, fixed in 4% PFA, permeabilized with 0.2% Triton-X/PBS, washed and blocked in SuperBlock™ T20 (PBS) Blocking Buffer (ThermoFisher cat# 37516). Primary antibody (diluted in SuperBlock™ T20 (PBS) Blocking Buffer) was incubated ON at 4°C, washed and incubated in secondary for 30 minutes, washed and mounted with a drop of mounting medium with DAPI (SouthernBiotech Cat#0100-20). Coverslips were sealed with nail polish and slide stored in dark at -20°C or +4°C. Primary antibodies include anti-GFP (aveslabs, cat# GFP-1010, 1:1000) and anti-S100 (abcam cat# ab41548,1:200).

### Imaging quantitation

Fluorescence imaging was performed using a Zeiss LSM 700 Confocal Microscope under identical acquisition settings for all groups within the same experiment to ensure comparability. Tissue sections were imaged at 40× or 63× magnification. For each animal, 3 anatomically matched fields were acquired per region of interest in a blinded manner when applicable. Quantitative analysis was performed using Fiji (ImageJ). All images were processed using the same analysis pipeline. Briefly, background fluorescence was subtracted using the “Subtract Background” function, and a consistent threshold was applied across all images. Mean fluorescence intensity and/or percentage of positive area was quantified for each field. The average of all fields per animal was used as a single biological replicate for statistical analysis.

### Statistical Analysis

Statistical analyses were performed in R. Biodistribution outcomes, including GFP, Luc, and ITR, were analyzed as continuous variables using mixed-design repeated-measures ANOVA to test whether vector distribution differed by AAV capsid, tissue, and/or their interaction. AAV capsid was modeled as the between-subjects factor, tissue as the within-subjects repeated-measures factor, and mouse ID as the subject identifier. Exon 17 expression was analyzed using the same repeated-measures framework. qPCR-derived dCt values were used for statistical analyses, and fold-change values relative to AAV-9 or PBS were calculated for graphical presentation. Model assumptions, including normality, homogeneity of variance, and sphericity, were assessed, with corrected p-values used when appropriate. Post-hoc comparisons were performed within tissue: Tukey-adjusted pairwise comparisons were used for GFP, Luc, and ITR biodistribution analyses, whereas Dunnett’s test was used for exon 17 expression to compare treatment groups against the PBS group. Statistical significance was defined as p < 0.05, and exact p-values, sample sizes, and detailed ANOVA/post-hoc results are reported in the corresponding figure legends.

## RESULTS

### ASO Development

Previously, we have shown that ASOs directed to *NF1* exon 17 exon splice enhancer (ESE) can be used to skip exon 17 *in vitro*^27^. We continued to develop our ASOs to increase the efficiency of exon skipping. Primarily, refined our ASO sequences and found that ASOs at +105 and +132 are the most potent *in vitro* when delivered as morpholino oligomers (PMOs) (Figure 1). Importantly, ASOs only recognize the human *NF1* exon 17 allele and not the mouse. The optimal PMOs have been aligned against the murine *Nf1* sequence and they have at least four mismatches. Human PMOs were evaluated in mouse cells, including Neuro-2a cells at 2 μM and mouse embryonic stem cells at 20 μM, and exon 17 skipping was absent in both (Supplementary Figure 1). The specificity of these PMOs for human *NF1* precludes testing *in vivo* exon-skipping efficacy in WT mice. This highlights the need for a mouse model carrying a humanized *NF1* exon 17, ideally designed to assess not only the efficiency of exon skipping but also the efficacy of skipping an exon containing a pathogenic variant in restoring *NF1* expression and activity.

**Figure 1:**
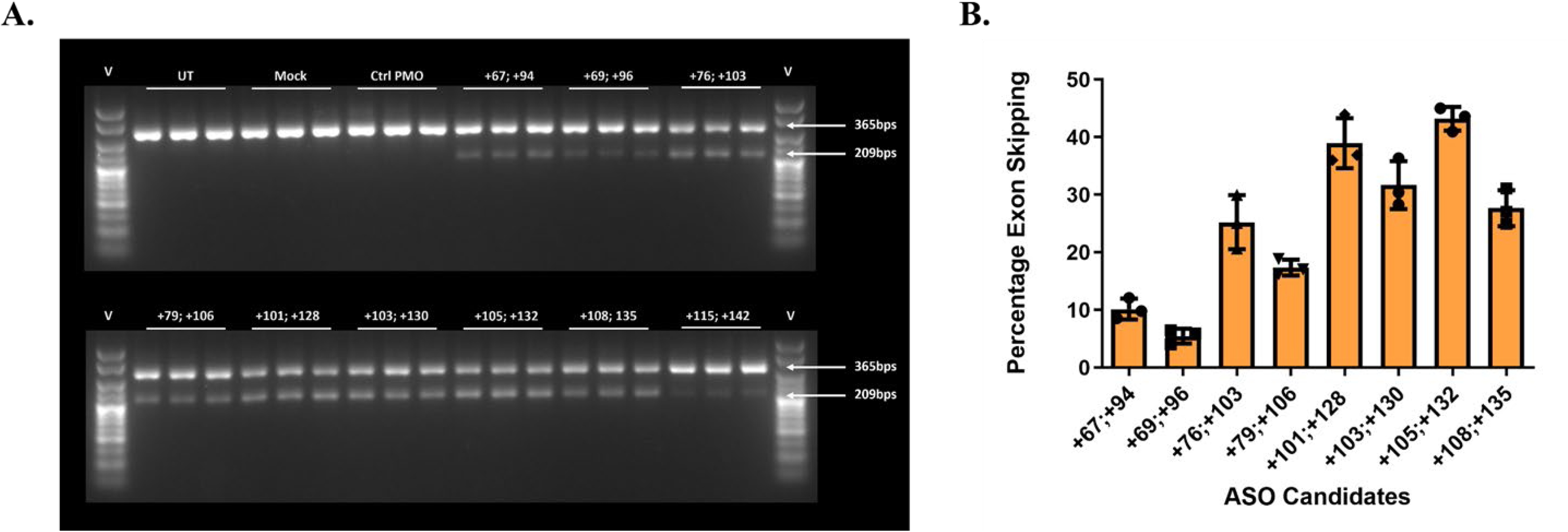
Antisense Oligonucleotide (ASO) Screening in HEK293T/A15; cultures engineered to harbor biallelic c.1885>A: G692R mutations in *NF1* Exon 17. A. Gel Electrophoresis and densitometric analysis of *NF1* Exon 17 skipping, for eight 28mer ASOs with a phosphorodiamidate morpholino oligo (PMO) chemistry. The ASOs were delivered at a 2µM dose using 6µM endoporter, to HEK293T/A15. After 48 hours of incubation, RNA was harvested, cDNA generated and subjected to nested PCR. The nested PCR yielded full-length and Exon 17 skipped amplicons, at 356 and 209bps, respectively. These were resolved on a 3% (w/v) agarose alongside bioline hyperladder V (25bps), permitting densitometric analysis using ImageJ. B. Histogram showing the percentage exon skipping quantified for the 28mer ASO candidates following densitometric analysis.

### Development of an AAV with U7-SnRNA approach and bioreporter

To enable analysis of biodistribution alongside examination of exon skipping efficacy of virally-delivered U7-SnRNA encoding the lead *NF1* exon 17 ASO construct, we have designed, cloned and validated a SFFV-driven transgene cassette compatible with multimodal imaging *in vivo* (Fluc) and post-mortem (EGFP) (Figure 2A). This construct was transfected into HEK293 cells at 1µg, 2µg, and 4µg doses and evaluated for GFP expression after 24 hours (Figure 2B). This data indicates dose response of GFP expression. Additionally, we evaluated the ability of the construct to impact *NF1* exon 17 skipping after 24 hours with increasing 1µg, 2µg and 4µg doses in HEK293T cells (Figure 2C and D). Densitometric analysis of PCR products indicates up to 25% exon skipping. AAV vectors carrying this expression cassette were packaged in AAV-9, AAV-F, and AAV-B1 capsids to explore tropism in different tissues through multiple delivery routes to allow usage in preclinical models. These serotypes were chosen on the basis that they had each been used previously for delivery to the brain by both ICV and IV routes in mice but no direct comparison between AAV-F and AAV-B1 has been documented.

**Figure 2:**
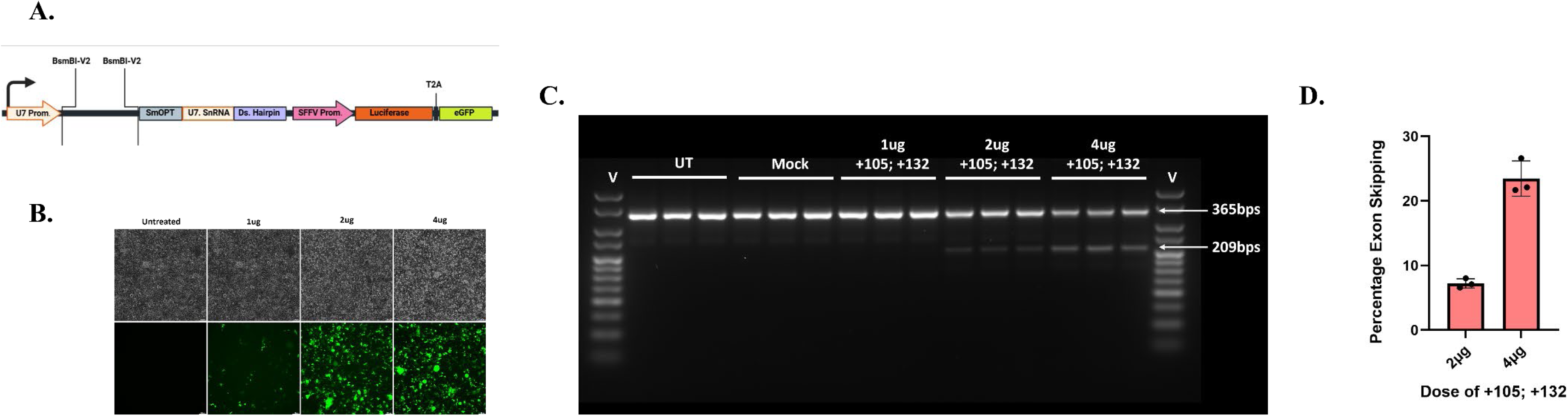
Validation of U7-SnRNA-SFFV-Luciferase-eGFP Constructs, in HEK293T cultures. A. Schematic highlighting the U7 construct sub-cloned into a pAAV backbone. Importantly, this comprises of a U7 promoter, Type IIS BsmB1-V2 restriction sites flanking a spacer sequence permitting the sub-cloning of ASO sequences, SmOPT a sequence optimized Sm-domain with high RNA binding affinity, U7 small nuclear RNA which can be directed by ASO sequences to interact with the spliceosome and a downstream hairpin to stabilize the structure A second transgene cassette comprising of an SFFV-promoter, Luciferase, a T2A linker and eGFP is also present to facilitate assessment of biodistribution. **B.** Brightfield and eGFP fluorescent microscopy, undertaken with 1µg, 2µg and 4µg doses of pAAV-U7-+105; +132-SnRNA-SFFV-Luciferase-eGFP. **C.** Gel Electrophoresis and densitometric analysis of *NF1* exon 17 skipping with increasing 1µg, 2µg and 4µg doses of pAAV- U7-+105; +132-SnRNA-SFFV-Luciferase-T2A-eGFP. The construct was transiently transfected into HEK293T using a 5:1 ratio of Viafect: DNA. After incubation for 48 hours, RNA was harvested, cDNA synthesized and nested to a nested PCR. The full-length and exon skipping amplicons 356 and 209bps respectively were resolved on a 3% (w/v) agarose alongside hyperladder V (25bps). Densitometric analysis was then undertaken using Image J. **D.** Histogram showing the percentage exon skipping quantified for the escalating dose of pAAV-U7-+105-SnRNA-SFFV-Luciferase-T2A-eGFP.

### Creation of humanized mouse model

In order to test human-specific ASOs preclinically, we created a mouse model with a humanized *NF1* exon 17 allele containing the patient-specific G629R pathogenic variant inserted 5’ of the mouse exon 17 using CRISPR/Cas9; termed hG629R mouse (Figure 3A). This mouse with the hG629R variant is expected to recapitulate the human mutation, leading to mis-splicing and a frameshift that will result in a pathogenic phenotype. Once germline, heterozygous matings were established and we determined that the allele is nullizygous lethal as Chi Square analysis of 37 pups born with 16 wildtype (WT) and 21 heterozygous (Het) and 0 homozygous (Hom) yields a two-tailed p-value of 0.0007. Brain tissue lysates from both WT and Het mice were evaluated at the RNA and protein level (Figure 3B). RNA transcripts from Het mice show both the wildtype and insertion alleles (259bp and 374bp, respectivel). Further, Het mice produce less than 50% neurofibromin protein in the brain as compared to WT mice (Figure 3 B, C).

**Figure 3.**
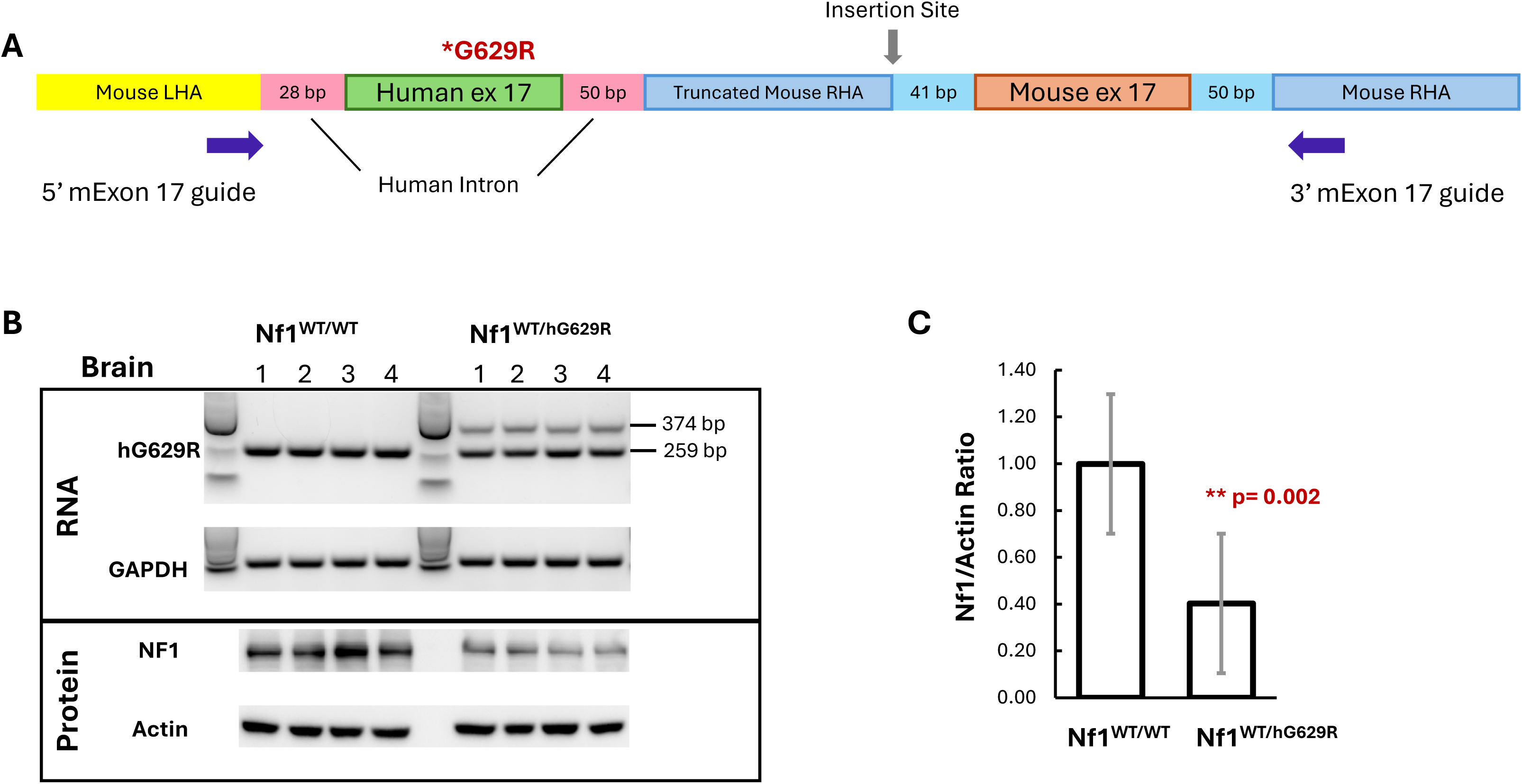
Generation and validation of the *Nf1* humanized exon 17 G629R mouse model. **A.** Cartoon depicting humanized exon 17 allele generated via CRISPR/Cas9. **B.** Representative RNA and protein analyses from *Nf1^WT/WT^* and *Nf1^WT/hG629R^* mice. RNA depicts WT allele (at 259 bp) in both WT and Het mice and both WT and insertion allele (374bp) in Het mice. GAPDH was used as a housekeeping control. Protein shows that Nf1 levels in Het mice are decreased in relation to WT mice. Actin used as a loading control. **C.** Quantification of neurofibromin protein levels normalized to actin in WT and Het mice. Data are shown as mean ± SEM; n = 4 mice per genotype. Statistical significance was determined by unpaired two-tailed t-test; **p = 0.002.**

### Capsid biodistribution comparison

To explore capsid tropism, we conducted a biodistribution study comparing them for ICV and IV administration routes. We also utilized this opportunity to standardize assessment methods for cross-comparison between tissues and with future planned studies. We utilized the Het hG629R mice (N=5 mice per group). Each mouse received PBS, AAV-9-U7-+105;+132-SFFV-Luc-eGFP, AAV-F-U7-+105;+132-SSFV-Luc-eGFP, or AAV-B1-U7-+105;+132 SFFV-Luc-eGFP. Mice were dosed IV at 5E13 vg/kg and ICV 1E13 vg/kg. Tissues were harvested 6 weeks post-administration and used to assess vector biodistribution and exon skipping at the DNA, RNA, protein, or histological level. The treatment and analysis scheme are depicted in Figure 4.

**Figure 4:**
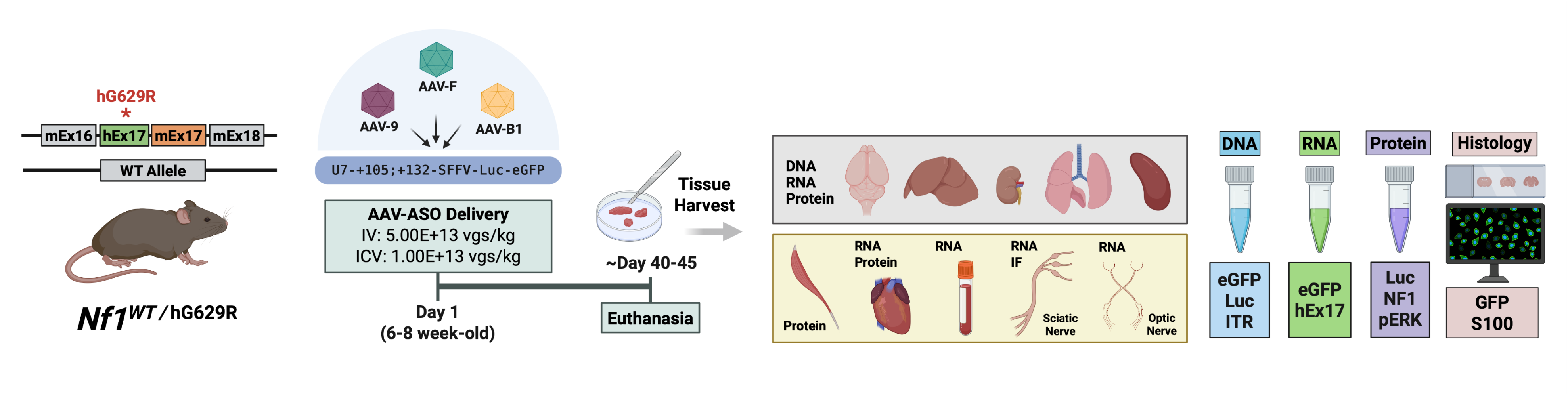
Cartoon depicting treatment scheme as well as the analysis of various indicated tissues. *Nf1* ^WT/hG629R^ mice harboring the human G629R point mutation in humanized exon 17 were injected on day 1 with AAV-ASO constructs (AAV-9, AAV-F, or AAV-B1 serotypes) carrying a U7+105;+132-SFFV-Luc-eGFP cassette via intravenous (IV; 5.00×10¹³ vgs/kg) or intracerebroventricular (ICV; 1.00×10¹³ vgs/kg) routes. Tissues were harvested at approximately day 40–45 post-injection following euthanasia. Indicated tissues were collected and analyzed at the DNA, RNA, and protein levels. Readouts included eGFP, Luciferase, and ITR at the DNA level by qPCR; hExon17 and eGFP RNA by RT-qPCR; Luciferase protein by luciferase activity assay; Nf1 and pERK protein by western blot; and GFP and S100 immunofluorescence in the sciatic nerve.

Interestingly, the highest AAV genome content per diploid cell was observed in the livers of mice injected ICV, with comparable levels across capsids (Figure 5A). In the brain, AAV-F was superior to both AAV-9 (p=0.006) and AAV-B1 (p=0.013). No significant differences were observed in any other tissue. Biodistribution analysis with ITR- and Luc-specific primers gave similar results (Supplementary Figure 2 A-B). Analysis of GFP mRNA levels revealed a similar relation between capsids in the brain (AAV-F > AAV-9 = AAV-B1) (Figure 5B). Interestingly, despite comparable vector genome content in peripheral tissues, there were significant differences in some peripheral tissues (kidney, spleen, and heart) where AAV-F and AAV-B1 expressed lower GFP mRNA levels than AAV-9 (Figure 5B). Moreover, mRNA expression levels in liver and brain were comparable for all capsids, despite the >10-fold differential in vector genome content between these tissues. Consistent with the mRNA expression data, luciferase activity was the highest in the brain for AAV-F across all tissues and capsids (Figure 5C). We also see significant increases in expression in muscle with AAV-9 over AAV-B1 (p=0.040) (Figure 5C). We found no evidence of exon skipping for most tissues, except in the sciatic nerve of the AAV-9 group (p<0.001) (Figure 5D). Together, these data indicate that AAV-F is the best capsid among those studied here for CNS gene transfer via an ICV delivery route, but that gene transfer efficiency may be too low to detect U7-driven exon skipping in the brain.

**Figure 5:**
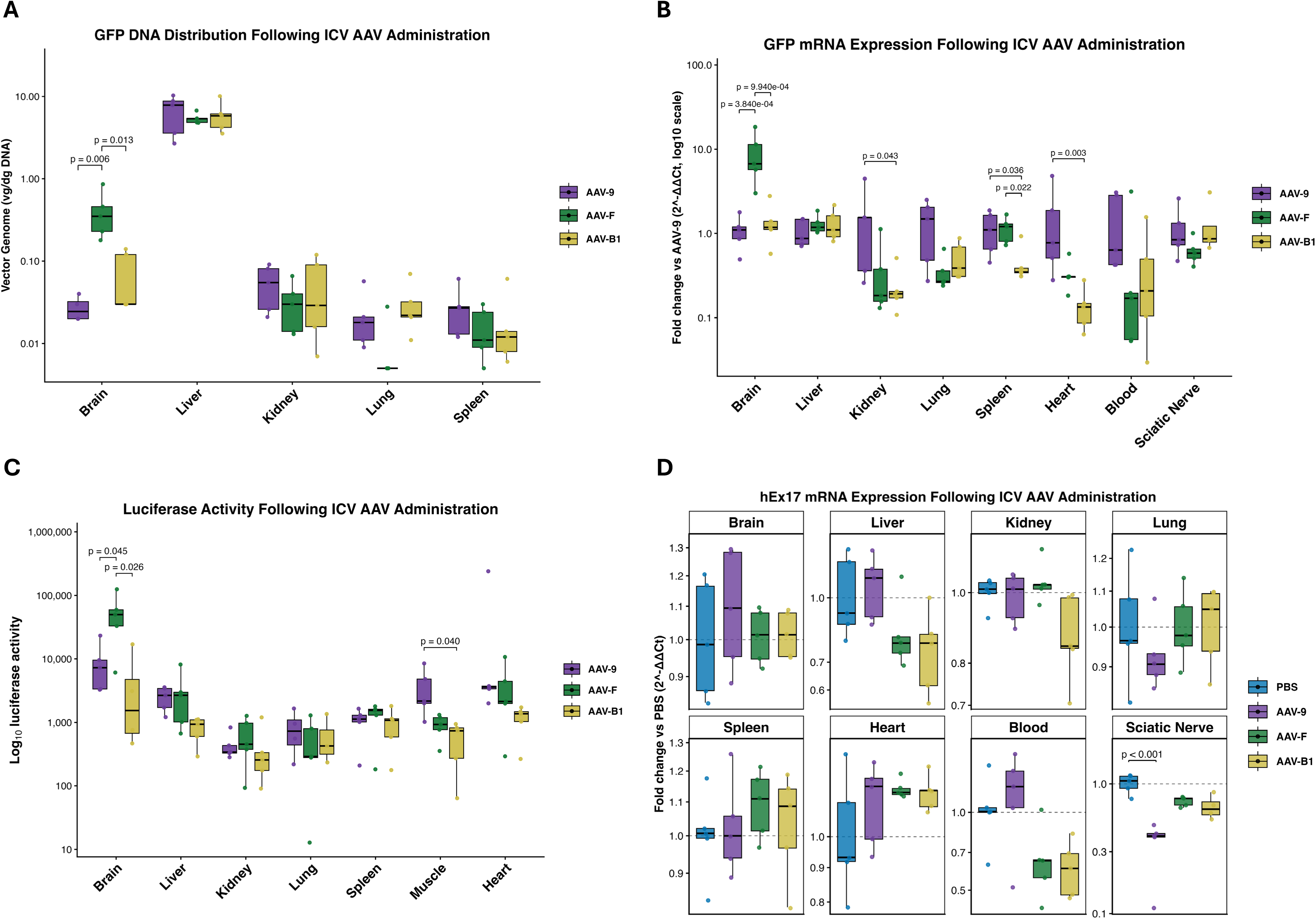
Capsid distribution and humanized exon 17 skipping following ICV AAV administration in mice. **A.** GFP DNA sequences were detected by qPCR in the indicated tissues following ICV administration of AAV-9, AAV-F, or AAV-B1 and are shown as vector genomes per diploid genome. **B.** Relative GFP mRNA levels measured by RT-qPCR in indicated tissues and are shown as fold change relative to AAV-9 using the 2^-ΔΔCt method. **C.** Luciferase enzymatic activity was measured and is shown as log-10 transformed in indicated tissues. **D.** Humanized exon 17 mRNA expression was quantified by RT-qPCR in the indicated tissues following treatment and is shown as fold change relative to PBS. Boxplots show the median and interquartile range, with whiskers extending to the most extreme values within 1.5× the interquartile range. Each point represents an individual mouse; n = 5 mice per treatment group. Mixed repeated-measures ANOVA was performed with AAV capsid as the between-subjects factor and tissue as the within-subject repeated-measures factor. Tukey’s multiple-comparison test was used for post hoc comparisons in panels **A–C**. Dunnett’s multiple-comparison test was used in panel **D** to compare treated groups against the PBS reference group within each tissue. Only significant comparisons are shown; adjusted p-values are indicated above brackets.

In the IV delivery study, as expected, liver had the highest AAV vector genome content, and AAV-F was significantly higher than AAV-9 (p=6.72e-05) and AAV-B1 (p=6.44e-05) in brain (Figure 6A and Supplementary Figure 3 A, B). Again, each assay marker (GFP, ITR, Luc) gives similar results for viral DNA transduction. Focusing on GFP sequences, we see that all capsids home to the liver the most (Figure 6A). Also, AAV-F is relatively de-targeted from other peripheral tissues compared with AAV-9 and AAV-B1, namely in the lung and spleen (Figure 6A). Analysis of mRNA expression levels showed the same profile, with AAV-F showing the highest expression levels in the brain (Figure 6B). No differences were apparent in other tissues. The luciferase activity in the brain was significantly higher for AAV-F compared to AAV-9 (p=0.011) and AAV-B1 (p=0.008), (Figure 6D). No other statistically significant differences are seen.

**Figure 6:**
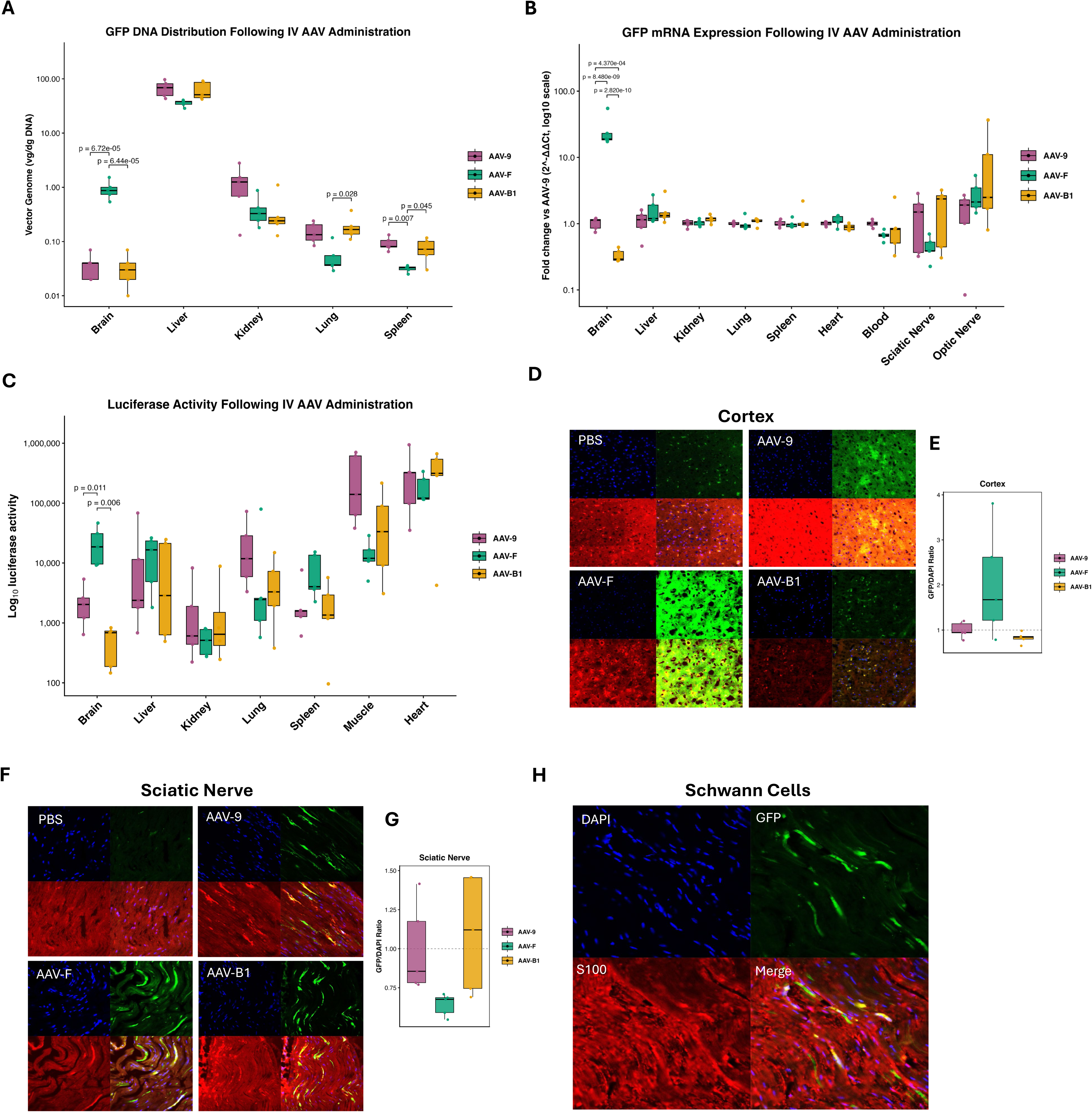
Capsid distribution following IV AAV administration in mice. **A.** GFP DNA sequences were detected by qPCR in the indicated tissues following IV administration of AAV-9, AAV-F, or AAV-B1 and are shown as vector genomes per diploid genome. **B.** Relative GFP mRNA levels were measured by RT-qPCR in indicated tissues and are shown as fold change relative to AAV-9 using the 2^-ΔΔCt method. **C.** Luciferase enzymatic activity was measured and is shown as log-10 transformed in indicated tissues. **D.** Representative fluorescence images showing GFP reporter expression in the cortex. **E.** Quantification of cortical GFP signal, expressed as the GFP/DAPI ratio, from three images per mouse. **F.** Representative fluorescence images showing GFP reporter expression in the sciatic nerve. **G.** Quantification of sciatic nerve GFP signal, expressed as the GFP/DAPI ratio, from three images per mouse. **H.** Representative sciatic nerve immunofluorescence images showing DAPI-stained nuclei in blue, GFP reporter signal in green, and the Schwann cell marker S100 in red. The merged image shows overlap between GFP reporter signal and S100-positive Schwann cells, appearing yellow. Boxplots show the median and interquartile range, with whiskers extending to the most extreme values within 1.5× the interquartile range. Each point represents an individual mouse; n = 5 mice per treatment group. Mixed repeated-measures ANOVA was performed in panels **A-C** with AAV capsid as the between-subjects factor and tissue as the within-subject repeated-measures factor. Tukey’s multiple-comparison test was used for post hoc comparisons. One-way ANOVA was performed in panels **E and G** followed by Tukey’s multiple-comparison test for post hoc comparisons. Significant comparisons, when present, are shown with adjusted p-values indicated above brackets.

In addition to the molecular assays, we evaluated GFP expression in histological tissue sections. We observed higher GFP expression in the cortex of mice in the AAV-F group, but quantitative analysis showed no statistically significant difference between capsids, possibly due to large variability across animals in the AAV-F group (Figure 6D, E). All capsids transduced cells in the sciatic nerve (Figure 6F, G). Image analysis showed that AAV-B1 may be slightly more efficient than AAV-9 and AAV-F, but there is substantial variability in transduction efficiency within the AAV-B1 group, so the difference was not statistically significant (Figure 6G). Regardless, we also show colocalization of AAV-F-delivered GFP with S100-positive Schwann cells, those most clinically relevant to neurofibroma development (Figure 6H).

As our end goal is to evaluate exon skipping in NF1-relevant tissues as a therapeutic strategy, we evaluated tissues for the presence of the humanized exon 17 allele. We see significant exon skipping in the liver with AAV-F (p=0.032) and AAV-B1 (p=0.009), in the sciatic nerve with AAV-B1 (p=0.002), and the optic nerve with all three capsids (Figure 7A). Other tissues showed no significant evidence of exon skipping. Since the brain is both relevant to NF1 and shows some evidence of exon skipping at the mRNA level, we looked for restoration of neurofibromin protein via Western blot and see that all capsids have efficacy relative to PBS-treated mice. The protein expressed holds functionality; pERK/ERK ratios are significantly reduced with both AAV-F and AAV-B1 in relation to both PBS and AAV-9 (Figure 7B, D). Further, we plotted the abundance of humanized exon 17 skipping as a function of pERK/ERK ratio and observed a correlation with a non-zero slope. We also note some clustering of groups (Figure 7E).

**Figure 7:**
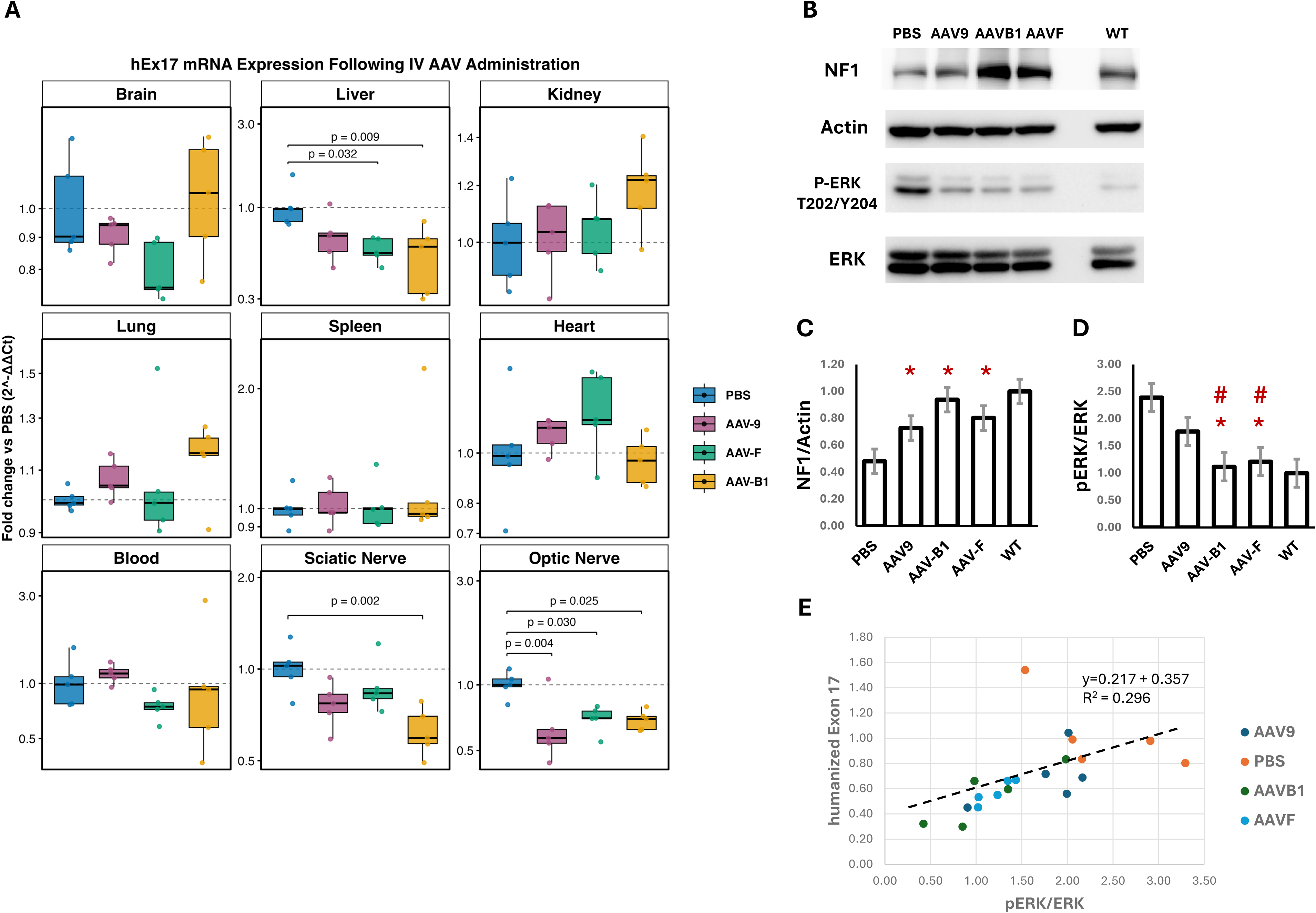
Humanized exon 17 skipping and neurofibromin protein restoration after IV delivery of capsids. **A.** Humanized exon 17 mRNA expression was quantified by RT-qPCR in the indicated tissues following treatment with the indicated capsids and is shown as fold change relative to PBS. Boxplots show the median and interquartile range, with whiskers extending to the most extreme values within 1.5× the interquartile range. Each point represents an individual mouse; n = 5 mice per treatment group. Mixed repeated-measures ANOVA was performed with AAV capsid as the between-subjects factor and tissue as the within-subject repeated-measures factor. Dunnett’s multiple-comparison test was used to compare treated groups against the PBS reference group within each tissue. Only significant comparisons are shown; adjusted p-values are indicated above brackets. **B**. Representative Western blot images of Nf1/actin and pERK/ERK in brain lysates from hG629R mice treated with indicated capsids. **C.** Quantitation (N=3 replicates; 5 mice each) of Nf1/actin levels in Het hG629R mice in relation to WT mice after IV capsid delivery. **D.** Quantitation (N=3 replicates; 5 mice each) of pERK/ERK levels in Het hG629R mice in relation to WT mice after IV capsid delivery. **E.** Correlation between humanized *NF1* exon 17 abundance and pERK/ERK ratios in brain lysates for each mouse in the indicated treatment groups. Statistical significance in panel **C-D** was determined using unpaired two-tailed t-tests. \**p < 0.05 compared with PBS; #p < 0.05 compared with AAV-9*.

**Figure 8:**
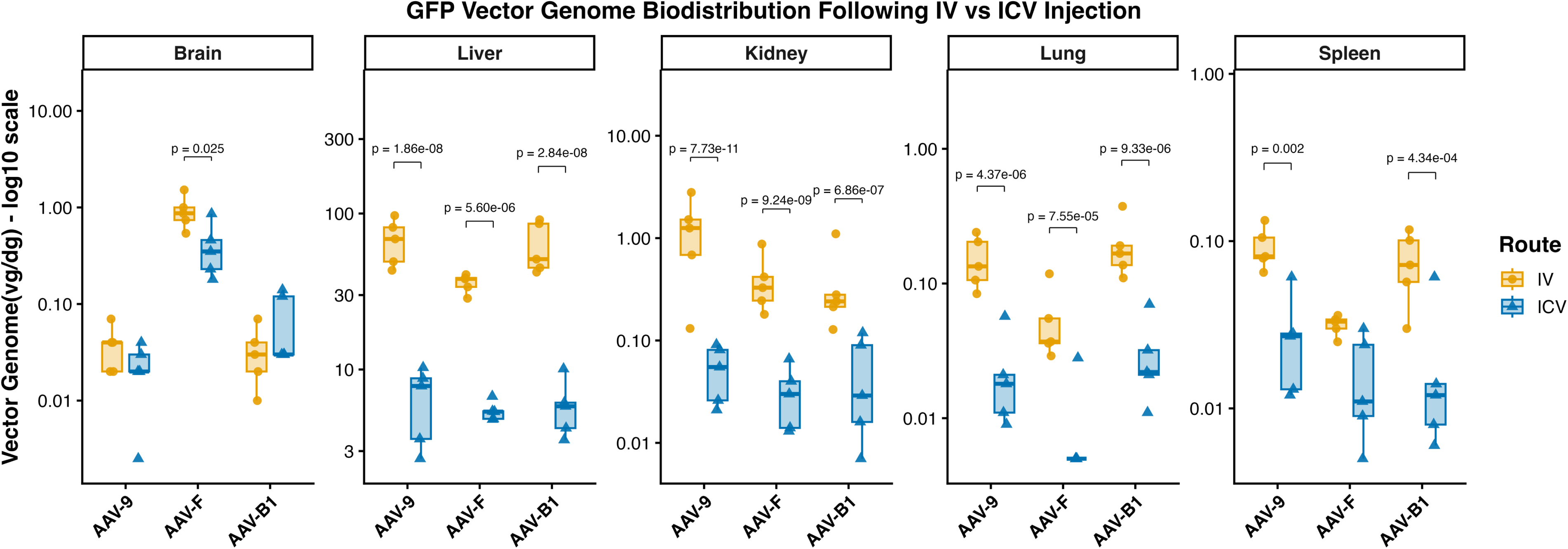
Route-dependent GFP vector genome biodistribution following AAV delivery. GFP vector genome copies per diploid genome were measured across tissues after IV or ICV injection of AAV-9, AAV-F, or AAV-B1 vectors. Each point represents an individual mouse tissue sample; boxplots show the median, interquartile range, and range. Log10-transformed values were analyzed using a linear mixed-effects model with Route, AAV capsid, Tissue, and their interactions as fixed effects, with Mouse ID included as a random intercept to account for repeated measurements across tissues from the same animal. IV versus ICV comparisons were performed within each AAV capsid and tissue using estimated marginal means with Holm correction. Exact Holm-adjusted p-values are shown. n = 5 mice per treatment group.

To evaluate differences between delivery routes, we can compare DNA transduction as it involves absolute quantitation (vector genomes/diploid genome). DNA transduction is highest in liver, regardless of route (Figures 5A, 6A and 8). Despite differences in dose (5E13 vg/kg IV and 1e13 vg/kg ICV), we anticipated that ICV delivery would result in more brain transduction than IV delivery; however, this is only significant for AAV-F (p=0.025). Further, ICV delivery results in roughly 10-fold less biodistribution to liver, kidney, lung, and spleen than IV delivery.

## DISCUSSION

The investigation presented here sought to address key translational barriers to the development of *NF1* Exon 17 skipping. We have presented selection of lead antisense sequence and vectorized this into an archetypal U7-SnRNA construct. By encoding the U7-SnRNA and Luciferase/eGFP reporters within a single AAV vector, we sought to examine *in vivo* biodistribution in AAV-9, AAV-B1 and AAV-F capsids and assess the feasibility of exon-skipping as a potential NF1 therapeutic intervention in tandem. Importantly, the humanized *Nf1* Exon 17 mouse generated permitted the screening of this advanced therapeutic. A comparative study of these three vectors has not been previously reported. This marks a step forward in AAV serotype validation for preclinical gene therapies targeting central and peripheral nervous system disease indications, such as NF1.

Screening of 28mer Phosphorodiamidate Morpholino Oligomers (PMOs) in HEK293T/A15 harboring biallelic G629R mutations in *NF1* Exon 17^27^ identified the lead candidate for targeted exon 17 skipping to be +105; +132. PMOs of 28 nucleotide length balance their biophysical properties, synthesis constraints and hold potential for conjugation to delivery moieties^29–31^. In addition to the AAV-U7-mediated exon skipping showcased here, it is anticipated NF1 patients might benefit therapeutically from alternative ASO modalities; chemical formulations would permit repeated administration and intratumoral injection. Interestingly, efforts to improve ASO transportation across the blood brain barrier (BBB) has been spearheaded by Spinraza, an ASO therapeutic for spinal muscular atrophy (SMA)^30^. In this instance, SMA targeting ASOs have been conjugated to PIP6a^32^ and ApoE peptides^33, 34^ and, a transferrin receptor targeting antibody ‘8D3’^30, 35^. Whilst such approaches improve BBB transcytosis, it is unknown whether the ASO will be delivered in sufficient quantities to achieve therapeutic benefit without toxicity. Importantly, however, PMO conjugates have the potential to be targeted to the CNS/PNS, addressing NF1 clinical manifestations, thus our optimal 28mer PMO described here is positioned to be exploited within this active area of research.

Here we packaged the optimal 28mer ASO as a U7-SnRNA expression cassette into a number of AAV serotypes. One of our primary goals was to assess the biodistribution achieved by the selected neurotropic capsids by the inclusion of the SFFV-Luciferse-T2A-eGFP cassette. This was assessed by assaying for vector transduction at the genomic level (qPCRs for vector ITR, GFP, Luciferase), expression of GFP and luciferase at the RNA (RT-qPCR) and protein levels (IHC for GFP and bioluminescence assay for luciferase), and quantification of targeted human *NF1* exon 17 skipping.

Assay results within individual studies show strong concordance between expression of bioreporters, but viral DNA transduction does not always predict bioreporter expression. As expected, levels of viral transduction markers GFP, ITR, and Luc are similar to each other within individual routes of delivery. Interestingly, with ICV delivery, highest transduction with all three serotypes is seen within the liver, but the highest eGFP mRNA expression and luciferase protein expression is seen within the brain. There is report of AAV genome silencing in the liver, which may explain this observation^36^. Levels of expression are highest with AAV-F. With the IV delivery route, we again see that viral transduction is highest in liver for all three serotypes, but that is not reflected in luciferase expression, which is highest in heart and muscle. Relative GFP mRNA expression still mirrors luciferase protein expression in the brain. This is supported by strong evidence of GFP protein expression by immunofluorescence in the cortex.

Interestingly, *NF1* exon skipping did not show concordance with transduction levels or bioreporter expression. This is exemplified in the liver with both ICV and IV routes of administration, where relatively low Luciferase/ eGFP expression is detected when considered in relation to the log-fold higher transduction displayed, but counterintuitively high exon skipping was observed. It is speculated that this dataset is largely attributed to the promoters driving the transgenes encoded; notably the U7 promoter supports sustained constitutive expression of U7-SnRNA transcripts^37^, whereas the SFFV is considered a ‘strong’ viral promoter^38^. In the liver, viral promoters typically have a short-lived spike of expression followed by a sharp decline, due to methylation and silencing^39^. Promoter dynamics could also go some way to explaining the apparent discrepancies between high Luciferase/ eGFP expression and low to modest exon skipping observed in the brain. The high expression of Luciferase/ eGFP from the SFFV promoter is good for determining biodistribution of vector but is not a reflective proxy of U7-SnRNA expression and its bioavailability to mediate exon skipping. It is possible that the cellular processing capacity limits the amount of U7 snRNA available for antisense delivery in a cell-type dependent manner.^40^ Alternatively, it may be that the expression profiles of the bioreporter relative to the SnRNA in some tissues is not proportional and expression of one may impact the other. Further influences including AAV-multiple transgene architecture^41^, and capsid-mediated histone modifications of the promoters or transgenes influencing expression cannot be ruled out^42^. If trying to achieve a more reflective proxy of therapeutic biodistribution for NF1 patients, future approaches should extend to *in situ* hybridization and/ or promoter selection more reflective of the constitutive, sustained expression characteristic of the U7 promoter, with an awareness of possible capsid influences.

Even with the consideration of AAV gene architecture and promoter discordance, it was surprising that no exon skipping was observed in the brain upon ICV administration of our AAV-F U7-SnRNA-reporter construct. Interestingly, however, evidence of exon skipping was observed in the blood, liver and sciatic nerve. This indicates that AAV-mediated exon skipping with a U7-SnRNA was functional. It is possible that the vector dosing may have also been too low to elicit higher levels of exon skipping. Two to three-fold higher vg/kg were reportedly required to achieve substantial therapeutic benefit of gene augmentation in neonatal SMA mice following CSF administration^43^. However, a recurring concern is that high AAV viral vector dosing is associated with significant morbidity and mortality in patient cohorts^44^. In consideration of this, future efforts to enhance exon skipping efficacy will focus upon vector genome design and capsid selection as opposed to increased dosing, as this will be more clinically relevant for NF1 patient cohorts. Alternatively, others in the field are using dual ICV and IV administration to achieve CNS therapeutic transgene expression in neonatal mice, which would indicate a potential benefit^45^. Notably for the work presented here, the AAV-F capsid appeared to retain some modest transduction benefits relative to AAV-9 in NHP models, particularly in the spinal cord and Schwann cells^13^.

Throughout our work AAV-F appeared to have improved tropism and bioreporter expression profiles to the brain irrespective of route of administration when compared to AAV-9 and AAV-B1. It also produced the highest levels of humanized exon 17 exclusion in the brain with IV treatment, though this was not significantly different to the other vector types. All three serotypes resulted in a significant increase in neurofibromin protein expression that acted to reduce pERK/ERK ratios towards wild-type levels. Notably AAV-F and AAV-B1 produced significantly greater reduction relative to AAV-9. We also see evidence for exon skipping after IV administration in the optic nerve at similar levels with all capsids. Exon skipping is also evident with IV treatment with all three vector types in the sciatic nerve, with AAV-B1 showing the highest efficacy. This differential potential of engineered AAV capsids to transduce central and peripheral nerves could have implications for NF1 patient cohorts, depending upon CNS/PNS tumor distribution. Clinically it could be argued that a systemic IV administration could hold greater translational value for NF1 patients given the multi-system manifestations of the disease. The 4-5% neurofibromin protein restoration reported recently to be enough to provide significant reduction in Ras-signaling^46^ is encouraging but significant questions remain as to whether this is the minimal level required in all tissue types and if this will be sufficient in later stages of disease progression. As further information related to this becomes available it will certainly change the NF1 advanced therapeutic landscape. The work reported here showed amelioration of aberrant Ras signaling, however, further investigations examining functional and physiological impacts in tumors would be required.

Importantly, for further translational relevance to NF1 patients, advanced therapies need to identify AAV capsids able to mediate BBB-transcytosis, as showcased by the vectors serotypes discussed so far, but also achieve expression of transgenes in Schwann cell lineages in the PNS. In addition, the use of surrogate capsids for translational models with the outlook to transfer to ‘human’ specific capsids, or development of AAV capsids with cross-species efficacy will be required. It is important to note that the AAV-F capsid engages a rodent specific receptor (Ly6c1)^17^ on the brain endothelium (akin to PHP.B engaging Ly6a^47^), thus the enhanced BBB crossing in mice will not translate to humans. Thus, in this study via the systemic route, AAV-F serves as a model AAV capsid to assess the effects of high transgene expression efficiency constructs of interest. That said, its ∼10-fold increase in transgene expression vs. AAV-9 via ICV delivery may be worth examining in large animal models, given it does not rely on Ly6c1 receptor engagement for crossing the brain endothelium for its properties via this route.

Since the commencement of this study additional AAV serotypes have been developed including SC3 and SC4^48^ and SCHW1 and SCHW2^49^. Crucially, SC3 and SC4 have been shown to have a 2.1-fold increased transduction of pNFs and a 6-fold increased transduction in the sciatic nerve compared to AAV-9 in mouse models, respectively. Furthermore, SCHW2 showed a 2.5 improved specificity towards Schwann cells *in vivo* in mice relative to AAV1 and a 57-fold reduction in targeting of the liver. Carneiro et al. developed two human Schwann cell-specific capsids C1 and C6^49^ and another group developed AAV-K55 which was designed to target human ST88-14 MPNST xenograft tissues in mice^11^. AAV-K55 demonstrated marked tropism to NF1-related neurofibroma, MPNST and glioma in xenograft mouse models, with greatly reduced liver transduction, and achieved significant therapeutic efficacies in treating NF1 ST88-14 xenograft tumors. There was however limited brain penetration and no detectable transduction of the sciatic and optic nerves in mice.

The current investigation vectorized ASOs into an archetypal U7-SnRNA construct into AAV capsids engineered for neurotropism. It is important to note that ASOs and U7-SnRNA constructs achieve modulation of splicing via distinct mechanisms. ASOs Watson-Crick base pair with pre-mRNA to steric block exonic splicing enhancer motifs and other regulatory elements in the pre-mRNA. In contrast the U7 encoding-SnRNA also base pair but express small nuclear Ribonucleoproteins (snRNPs), that recruit or manipulate splicing machinery. The U7-SnRNA construct used in this investigation, was based on that described in 2004 to achieve *DMD* exon skipping^28^. Unsurprisingly, significant sequence engineering to the canonical U7 RNA ‘chassis’ has been undertaken since the conception of this therapeutic intervention. Two examples with striking increases in exon skipping efficacy include fusion of the U7-SnRNA to a splicing silencer^50^ and the mutagenesis of U7-SmOPT to support *ADAR* RNA editing as a means of exon skipping^51^. These studies show promising increases in efficacy that could be exploited for the NF1-related exon skipping approach showcased.

Distinct from the direct engineering of the U7-SnRNA architecture, the length of the ASO sequence can also be modified. Canonical PMOs typically vary in length between 25-30mer due to synthesis constraints; in contrast ASO sequences encoded within the U7-SnRNA conventionally vary between 20-45 nucleotides. However, research by Shimo et al. has indicated that increasing the ASO lengths to 45-75 and 75-110 nucleotides enhanced exon skipping of the *FAS* and *DMD* genes respectively^52^. Whilst in this study modest exon skipping was observed on systemic administration with a canonical U7-SnRNA cassette, future efforts will examine extended ASO sequences and molecular engineering of the U7-SnRNA scaffold to enhance *NF1* exon 17 skipping efficacy.

The research presented here was proof-of-principle in nature; however, it is encouraging to recognize the current unrealized potential of the U7-SnRNA system presented and the significant opportunities to enhance exon skipping efficacy. Examination of therapeutic efficacy for NF1 in relevant mouse models with evaluation of skipping at the tumor level is required. Refinement of the U7-construct and ASO designs combined with engineered AAV capsids with cross-species efficacy could permit clinical translation and provide a mutation specific approach with relevance to NF1 patient cohorts.

## Acknowledgements

UAB Transgenic and Genetically Engineered Models Core; Gilbert Family Foundation’s Gene Therapy Initiative grant to Wallis and Popplewell.

## Authorship contributions

Conceptualization: LP, DW, RAK; Methodology: LP, DW, RAK, MM, CAM, HL, XZ, KRS; Providing Reagents: CAM, MSE; Investigation: MM, XZ, HL, EW; Validation: MM, XZ, HL, KRS, EW; Data Curation: MM, XZ, HL, KRS, TV, EW; Formal analysis: MM, KRS, TV, DW, LP, EW; Writing - Original Draft, DW, MM, HL, XZ, KRS, EW, LP; Writing - Review & Editing: DW, MM, LP, RAK, MSE, CAM, TV; Funding acquisition: DW, LP, RAK; Supervision: DW, LP, RAK; Project administration, DW, LP.

## Patents

Work related to this manuscript is covered by patent application 17/762,355: Methods of treatment of Neurofibromatosis Type I (NF1) and NF-1 Mediated Conditions and Compositions for Use in Such Methods.

---

**Supplementary Figure 1.**
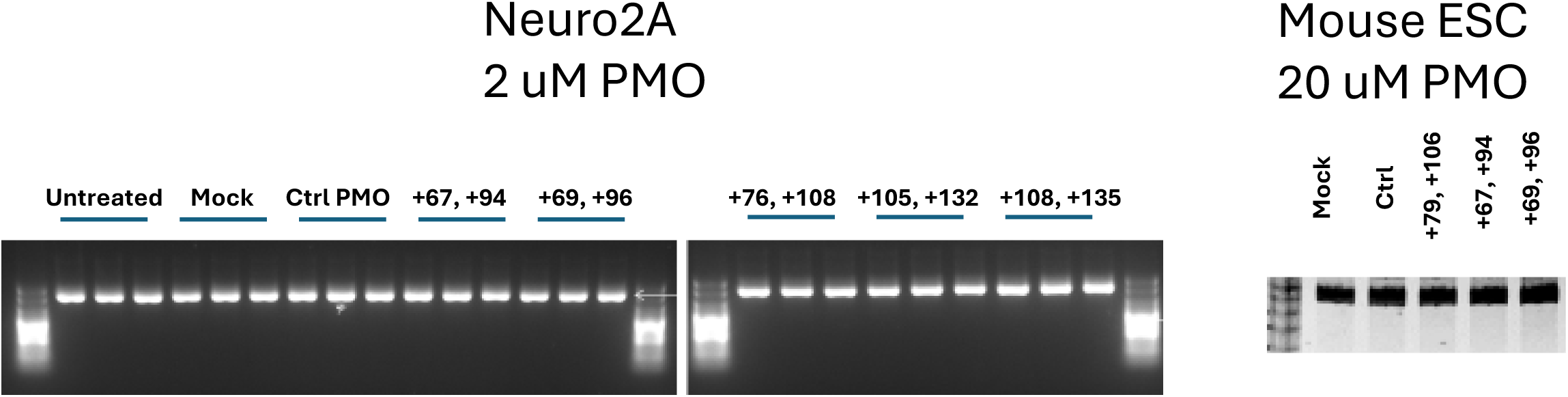
ASOs are specific to the human *NF1* exon 17 allele as no exon skipping is seen in mouse cells. Gel Electrophoresis analysis of *Nf1* Exon 17 skipping, for various indicated human PMOs that were evaluated in mouse cell lines including Neuro-2a cultures at 2 uM and in murine embryonic stem cells at 20 uM for exon skipping. After 48 hours of incubation, RNA was harvested, cDNA generated and subjected to PCR and resolved on a 3% (w/v) agarose,

**Supplementary Figure 2:**
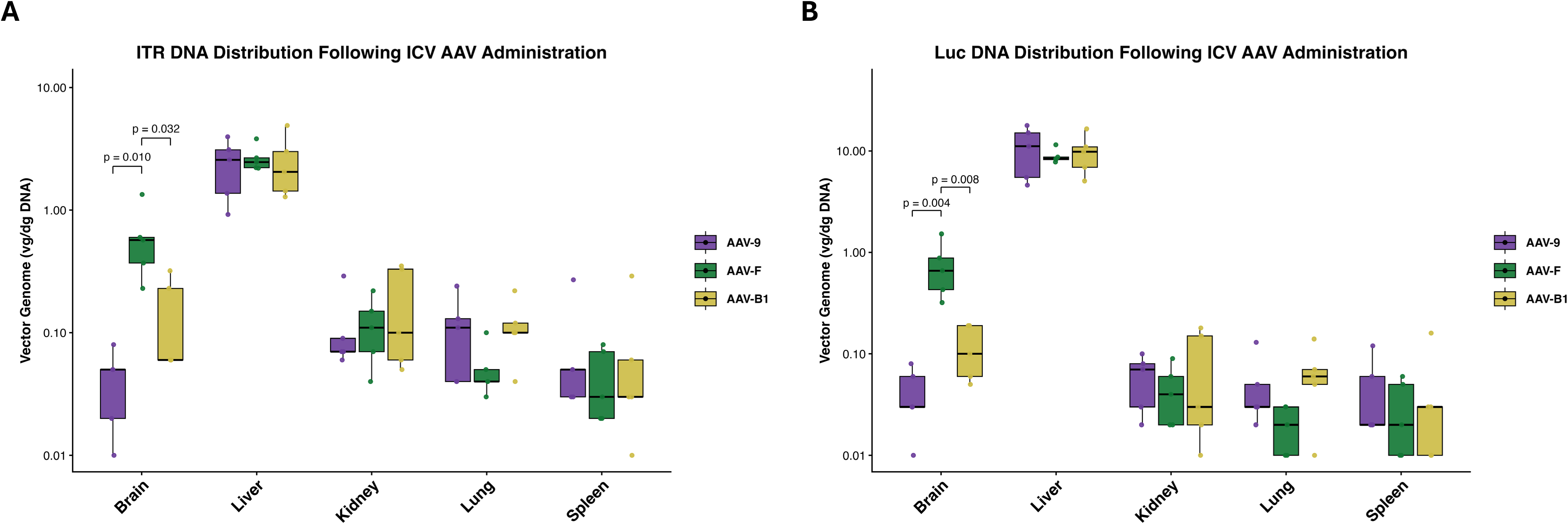
Capsid distribution following ICV AAV administration in mice. **A.** ITR, and **B.** Luciferase DNA sequences were detected by qPCR in the indicated tissues following ICV administration of AAV-9, AAV-F, or AAV-B1 and are shown as vector genomes per diploid genome. Boxplots show the median and interquartile range, with whiskers extending to the most extreme values within 1.5× the interquartile range. Each point represents an individual mouse; n = 5 mice per treatment group. Mixed repeated-measures ANOVA was performed with AAV capsid as the between-subjects factor and tissue as the within-subject repeated-measures factor. Tukey’s multiple-comparison test was used for post hoc comparisons. Only significant comparisons are shown; adjusted p-values are indicated above brackets.

**Supplementary Figure 3:**
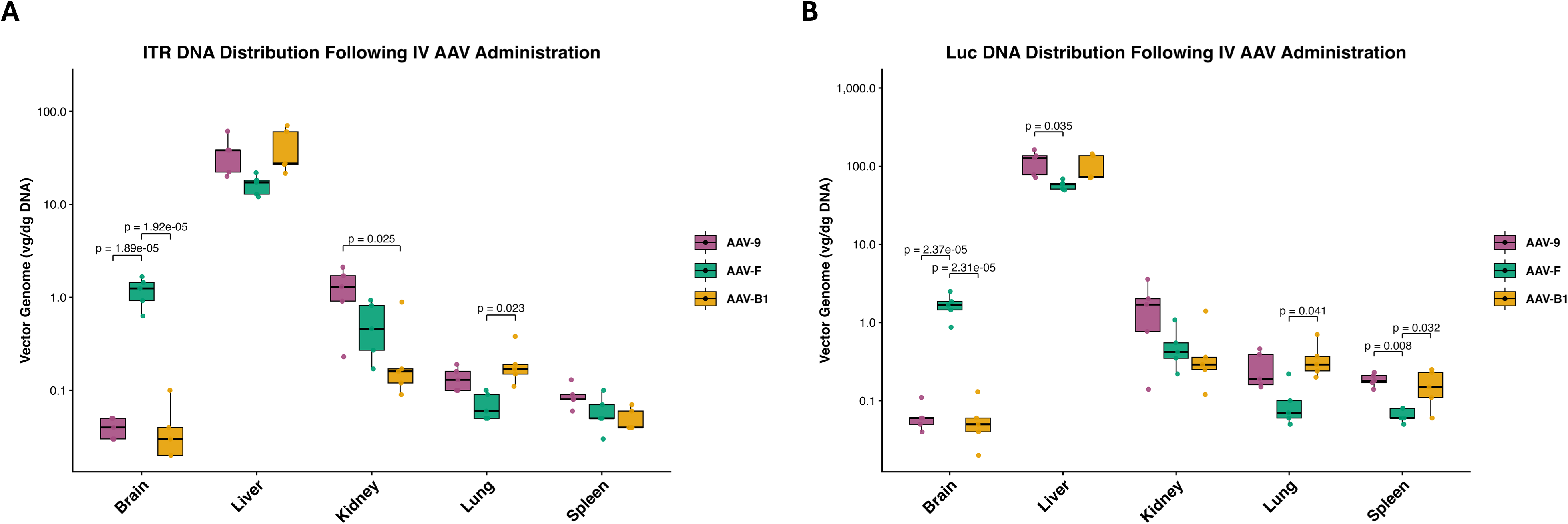
Capsid distribution following IV AAV administration in mice. **A.** ITR, and **B.** Luciferase DNA sequences were detected by qPCR in the indicated tissues following IV administration of AAV-9, AAV-F, or AAV-B1 and are shown as vector genomes per diploid genome. Boxplots show the median and interquartile range, with whiskers extending to the most extreme values within 1.5× the interquartile range. Each point represents an individual mouse; n = 5 mice per treatment group. Mixed repeated-measures ANOVA was performed with AAV capsid as the between-subjects factor and tissue as the within-subject repeated-measures factor. Tukey’s multiple-comparison test was used for post hoc comparisons. Only significant comparisons are shown; adjusted p-values are indicated above brackets.

## eTOC synopsis

We advanced *in vivo NF1* exon 17 skipping by optimizing antisense oligonucleotides, creating a humanized mouse model, and developing AAV-U7-snRNA delivery. A biosensor packaged in AAV-9, AAV-F, and AAV-B1 revealed tissue-specific transduction. This is the first report of NF1 exon skipping efficacy *in vivo* with the restoration of functional NF1.

